# Humans progressively feel agency over events triggered before they act

**DOI:** 10.1101/2023.12.01.569449

**Authors:** Marcus Toma, Jérémie Mattout, Romain Quentin, Fayed Rassoulou, Alice Gautier, Emmanuel Maby, Marine Vernet

**Affiliations:** IMPACT Team, Lyon Neuroscience Research Center (CRNL), CNRS (UMR 5292), INSERM (U1028), University Claude Bernard Lyon (UCBL), Lyon, France; COPHY Team, Lyon Neuroscience Research Center (CRNL), CNRS (UMR 5292), INSERM (U1028), University Claude Bernard Lyon (UCBL), Lyon, France; EDUWELL Team, Lyon Neuroscience Research Center (CRNL), CNRS (UMR 5292), INSERM (U1028), University Claude Bernard Lyon (UCBL), Lyon, France

## Abstract

Artificial Intelligence (AI) has become increasingly efficient in anticipating our behavior. Will this impact, in the near future, how much we feel control over events generated with AI-assistance? This would in turn influence our decisions, actions, mental health and sense of responsibility. In everyday life, our sense of agency over events occurring at various delays after our actions has adapted to accommodate these delays. Here, we investigate whether our sense of agency can also adapt to an unusual situation where a consequence precedes an action. We used an online game in which players aimed to beat the computer at finding and clicking on a target to trigger an animation, while, in fact, an algorithm triggered the animation before the players’ click. The animation was not randomly controlled by the algorithm, but rather based on the history of the players’ past movements and on the beginning of their current movement. We used correlational, machine learning decoding, and modelling approaches to capture how players compute their reported sense of agency over the animation. We found evidence that, in less than an hour, players implicitly learned that the timing of the animation was related to their own actions, and adapted their sense of agency accordingly. Such findings may help us to anticipate how humans will integrate AI-assistance to guide their behavior.

## Introduction

Artificial Intelligence (AI) has become increasingly efficient in nudging and even anticipating our actions. To name a few, online shopping recommendations or social media content selection are known to influence our choices ^1–3^, while smart assistants, autocomplete modes, and predictive text generation based on individual past history are gradually becoming more accurate in predicting our thoughts and actions to the extent that we do not need to complete them anymore ^4,5^, generating uncertainty about the feeling of control of end-users ^6^. These AI outcomes generally share two characteristics: they occur earlier than what we would have produced without AI assistance, yet they are not independent from us: they are based on our past actions and the start of a current action. How much can we feel in control of the produced output? Does this feeling evolve as we become accustomed to the assistance?

Cognitive neuroscience has widely explored such subjective feeling of control over the outcomes of our actions, which has been called the “external” sense of agency (in opposition to a “body” sense of agency that would refer to feeling at the origin of our body movements ^7^), but is most often simply referred to as the sense of agency (SoA) ^8^. Such SoA is central to the fluency of action (i.e., the sense of ease with which an action is selected), responsibility and wellbeing in humans ^9^. It can be degraded in neuropsychiatric patients suffering from depression or schizophrenia ^10,11^. The SoA is believed to derive from both preparatory motor signals, and from a strong match between predicted and actual sensory consequences of an action ^12^. More generally, the SoA would be built on integrating several factors, also including, for instance, emotion, goal achievement and effort, weighting them depending on how reliable a source of information is supposed to be (for reviews, see ^7,9^).

One crucial factor that can impact the SoA is the perceived causality between one’s action and its consequence, which should partly derive from the perceived temporality of events. However, how can we infer precise temporal judgment of internal and external events, given that the neural temporal processing differs greatly across sensory modalities? It has been proposed that temporal perception is based on adaptive recalibration of such timings from our daily experience: several minutes of an exposition to a fixed audiovisual delay are sufficient to shift the perceived time of equivalence between auditory and visual signals ^13^. Such flexibility in the temporal perception of events would explain why *a priori* causal beliefs and agency beliefs could strongly influence the perception of temporal events ^14–17^ to the point that events can be subjectively reordered to match the perceived causal relationship between the events ^18^.

Is the SoA as flexible as temporal perception? Shorter delays between an action and its consequence usually lead to higher SoA. Nevertheless, this relationship depends on the frequency of the situation encountered in everyday life: we are now used to computers that respond very quickly to our button presses but we are still tolerating longer delays for an elevator to arrive after we call it, and we must be sufficiently exposed to the delay of a new shower mixer to be able to regain a sense of control over the temperature it delivers. This adaptability can lead us to feel less agent when exposed to delays shorter than expected ^19^. It can even cause a subjective reversal of action and feedback sensation if the delay to which we are adapted is suddenly removed ^20,21^.

In comparison, the SoA when feedback truly precedes an action is much less studied. In the early 1990’s, Dennett described the “Grey Walter’s precognitive carousel” experiment that he heard about in a conference in the 1960’s ^22^. Patients with implanted electrodes within the motor cortex would press a button on a controller to change the slides from a carousel projector. However, the controller button was a dummy and the slides were actually changed based on motor signals from the patients’ brain. Patients reported that they were startled to see the slide projector anticipating their decisions by changing the slides before they could press the button, as if the carousel was reading their mind, and that they were worried that by pressing the button, the slides would change twice ^22^. Dennett later explained that this experience has never been published and has proven difficult to replicate ^23^, although brain-computer interfaces based on decoding motor preparation from EEG signal have been designed ^24,25^. Yet, his interpretation about such potential results remained the same: the actual delay does not matter, because there is no single, fixed timeline that would allow one to feel like an agent: this feeling instead arises based on our expectations built over our past experience. In everyday life, we are rarely exposed to feedback preceding our action, except in well-delimited cases of the AI assisted actions we introduced earlier. According to the temporal order of events, having a strong SoA in such cases would be an illusion. Nevertheless, because the outcome is actually based on our past actions and the current action’s opening, it would not be. Given the flexibility of the SoA to a variety of timings, in the present study, we hypothesized that (1) outcome preceding action would initially lead to a low SoA and (2) a long enough exposure to this situation would allow us to adapt and regain agency over the early outcomes. To test these hypotheses, we designed an online experiment presented in the form of a game to be played on a computer. Our goal was to trigger an effect (a visual animation displayed on the screen) consistently before players’ action (a click on the left button of the computer mouse), and then to measure whether the players felt like they were at the origin of the effect or not.

## Results

To measure whether humans could feel agent over events occurring before their actions, we adopted the following strategy: (1) Players believed they were playing against the computer to find a visual target in a grid of distractors and click on it as fast as possible, before the computer. However, the time they spent finding the target during the visual search did not actually matter to the game; only their movement to the target *per se* was used by the algorithm to manipulate the timings of the visual animation perceived by the players. (2) Players were trained, without knowing it, to make their movements as predictable as possible. Therefore, they were not allowed to move the cursor before finding the target, which they indicated by pressing the spacebar. Moreover, after pressing the spacebar, they had to move the cursor as accurately and as fast as possible towards the target and click on it, otherwise, they received a warning and the current trial was started again. (3) An algorithm was trained on past movements to predict, on the current trial, when players would click on the target, and to start the visual animation shortly (∼80 ms) before. (4) Players then answered whether or not they thought that they “won”, i.e., that they clicked on the target before the computer to trigger the animation, which was taken as a measure of their SoA over the animation. (5) Trials were repeated (during ∼40 min) to test whether the SoA would evolve when players are consistently exposed to an effect preceding their actions.

### Correlational approach: players progressively increased their sense of agency over events occurring before their actions

All analyses were performed only on trials for which the computer’s virtual click (i.e., the animation) started before the players’ click (86 ± 11 % of the trials, with an average anticipation of 78 ± 21 ms). For each Block of trials, the uncorrected local SoA was calculated as the averaged SoA across trials (i.e., the ratio of trials in which the players answered they clicked first divided by all the trials). This uncorrected local SoA significantly increased across Blocks (linear fit coefficient=0.02 ± 0.03; significantly above 0: Wilcoxon t=639; p<10^−4^; effect size r_rb_=0.79, **Fig. 1 and 2A**) and the correlation coefficient between local SoA and Blocks was significantly positive (R=0.49 ± 0.48; significantly above 0: Wilcoxon t=650; p<10^−4^; effect size r_rb_=0.78).

**Figure 1:**
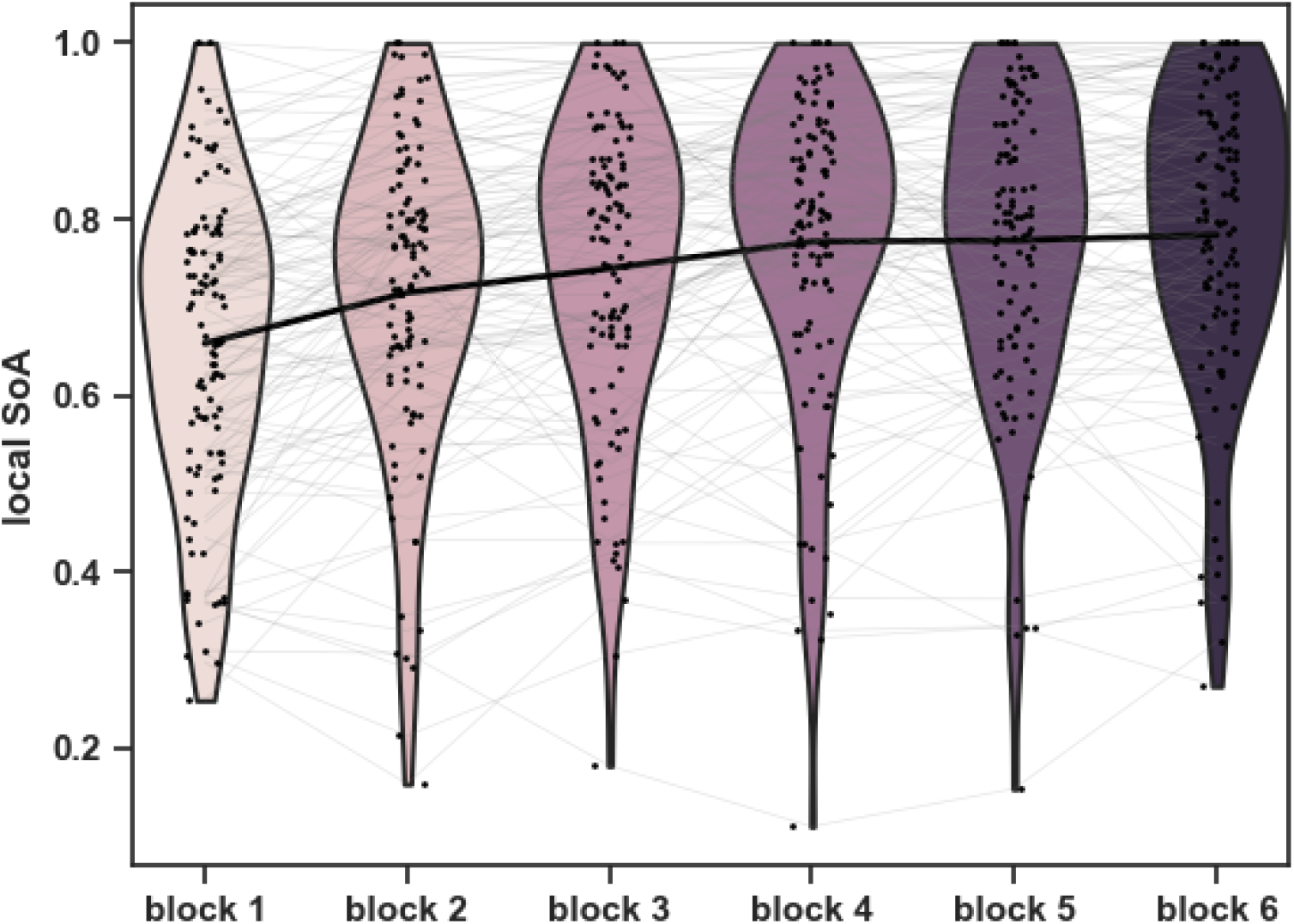
Increase of local SoA through the course of the game. The average local SoA was calculated for each block of trials as the ratio of trials in which the players answered they clicked first divided by all the trials in which the virtual click occurred before the human click. The violins represent the distribution of the local SoA for each block. Each dot and connecting thin line correspond to an individual player. The thick line represents the mean across players. Note the local SoA increase across the course of the game (linear fit coefficient=0.02 ± 0.03; significantly above 0: Wilcoxon t=639; p<10^−4^; effect size r_rb_=0.79).

**Figure 2.**
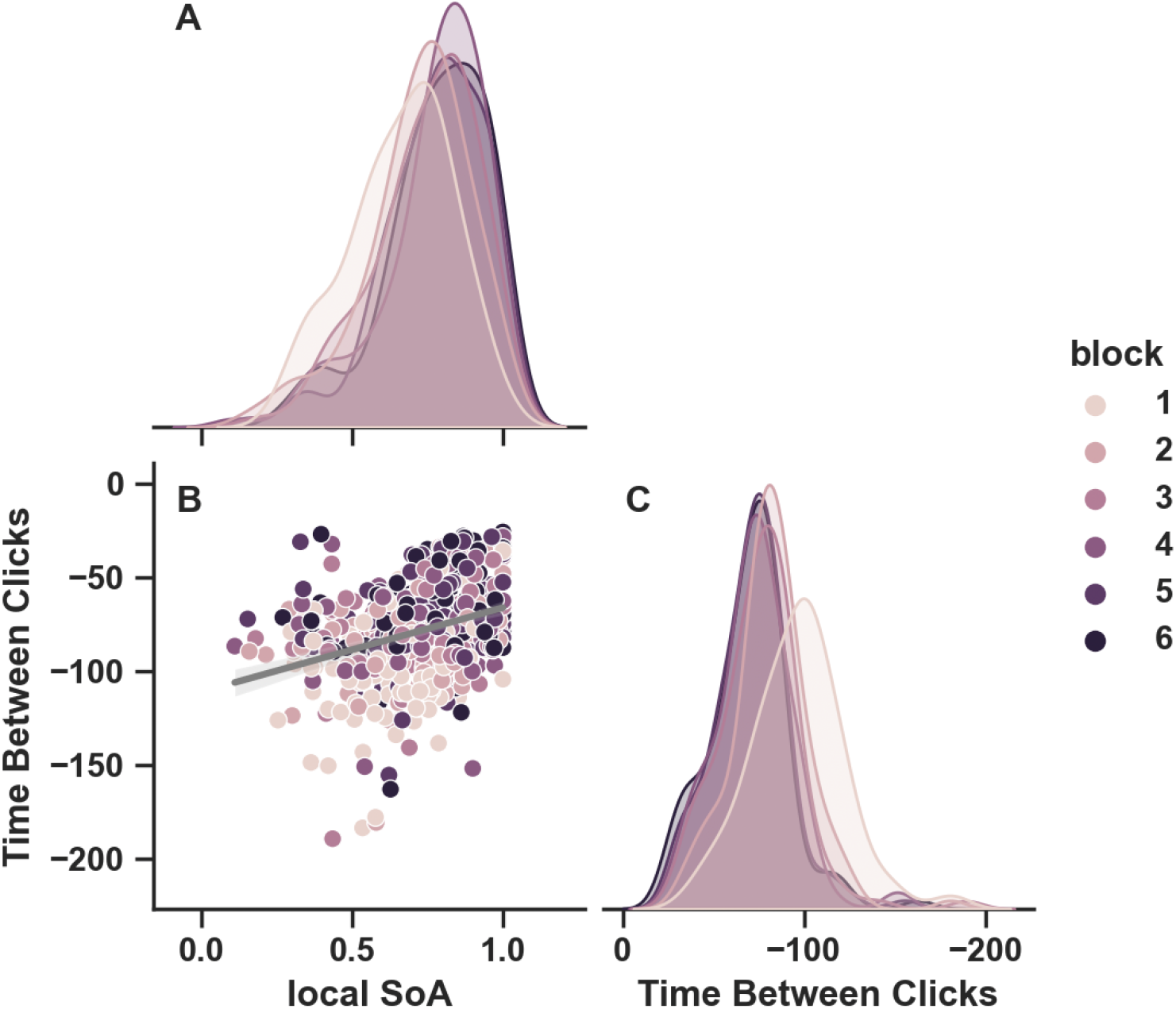
Distribution of the local SoA and of the Time Between Clicks for all Blocks and all players. **A**. Distribution of the local SoA for each block. The shift of the distribution illustrates the increase of local SoA through the course of the game. **B**. Correlation between the local SoA and Time Between Clicks. Each dot represents the average value for one participant and one block. The earlier the virtual click relative to the human click, i.e. the larger and more negative the Time Between Clicks, the lower the SoA (R=0.58 ± 0.40; significantly above 0: Wilcoxon t=225; p<10^−4^; effect size r_rb_=0.92). **C**. Distribution of the Time Between Clicks for each block. The shift of the distribution illustrates that through the course of the game, the virtual click happened closer and closer to the human click.

However, the precision of the algorithm also increased during the experiment, leading the Time Between Clicks (i.e., human click time minus virtual click time, averaged per Block) to become smaller **(Fig. 2C)**. As expected, the earlier the virtual click (relative to the human click), the lower the local SoA (**Fig. 2B**, on average R=0.58 ± 0.40; significantly above 0: Wilcoxon t=225; p<10^−4^; effect size r_rb_=0.92). Such correlation between Time Between Clicks and local SoA means that the players were appropriately playing the game and answering the SoA questions. To control whether the SoA increased independently of the decrease of the Time Between Clicks, the partial correlation between local SoA and Blocks taking into account the Time Between Clicks was computed. For this, local SoA was replaced by the residuals of the correlation between local SoA and Time Between Clicks, and Blocks were replaced by the residuals of the correlation between Time Between Clicks and Blocks. This corrected local SoA significantly increased across corrected Blocks (linear fit coefficient=0.01 ± 0.03; significantly above 0: Wilcoxon t=2240; p=0.0221; effect size r_rb_=0.25) and the correlation coefficient between residuals was significantly positive (R=0.15 ± 0.55; significantly above 0: Wilcoxon t=2080; p=0.0056, effect size r_rb_=0.31, **Fig. 3**). This demonstrates that players adapted to the game: the more they practiced the game, the more they felt they triggered the animation even though the animation started before their click. This result reveals that humans can quickly adapt their sense of agency to unusual situations, in which consequences of their actions precedes their actions.

**Figure 3.**
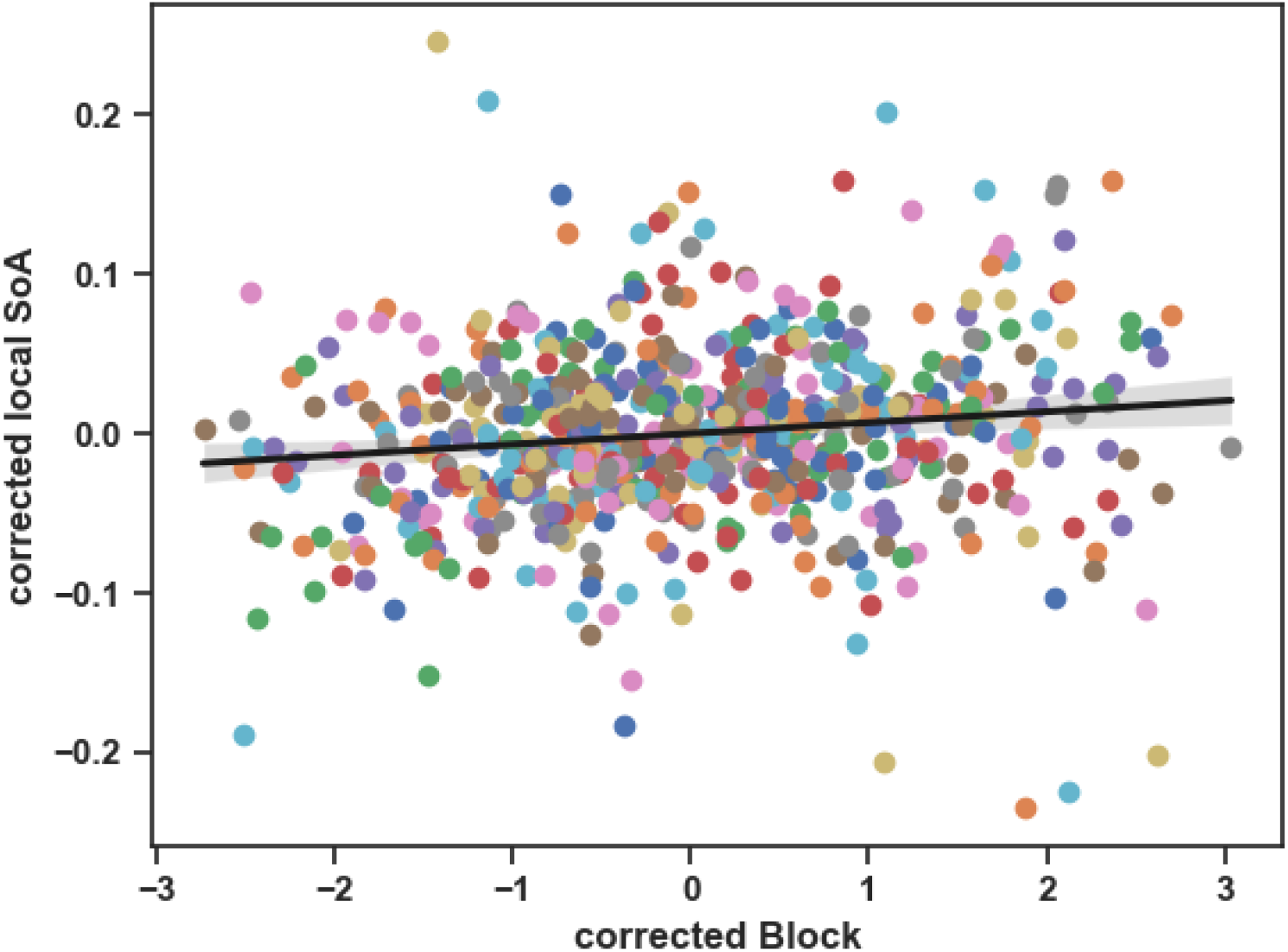
Correlation between corrected local SoA and corrected Block. The corrected local SoA corresponds to the residuals of the correlation between local SoA and Time Between Clicks and the corrected Block corresponds to the residuals of the correlation between Block and Time Between Clicks. Each color represents one player. The correlation between corrected local SoA and corrected Block was significant (R=0.15 ± 0.55; significantly above 0: Wilcoxon t=2080; p=0.0056, effect size r_rb_=0.31, not shown here for individual players for visibility). Taking all participants together, the correlation was also significant (R = 0.14; p = 0.0004; CI = [0.06, 0.21]).

Perfectly objective and accurate players should always report that the computer won. This was not the case: on average they answered that they “won” (i.e., that they provoked the start of the animation) 74 ± 15% of the time. Notwithstanding this mismatch, the above results demonstrated that players integrated objective information such as the Time Between Clicks to build-up their local SoA (with longer Time Between Clicks leading to lower SoA). It also demonstrated that the time spent playing the game independently increased their local SoA. Below, decoding and modelling approaches will confirm these results while exploring whether players used additional sources of information, such as their performance or other feedback from the game, to compute their local SoA.

### Decoding approach: the time spent in the game is predictive of the sense of agency

Machine learning models using Logistic Regression to predict the local SoA were applied on all possible combinations of the following parameters: Time Between Clicks, Blocks, Search Time (time between the start of the trial and the spacebar press), Waiting Time (time between the spacebar press and the beginning of cursor movement), Acceleration Time (time to peak speed after cursor movement onset), Peak Speed, Distance at Start (cursor distance to the target at the start of the trial), Distance at Human Click (cursor distance to the target when the player clicked), Distance at Virtual Click (cursor distance to the target when the animation was triggered), Human Click Time and Virtual Click Time (see **Supp. Fig. 1** for details on the evolution of these parameters during the course of the game). The maximum decoding performance was obtained using Blocks, Peak Speed, Distance at Virtual Click and Time Between Clicks and had an Area Under the Curve (AUC) of 0.86 ± 0.08 (significantly above chance (0.5): Wilcoxon t=0.00; p<10^−4^; effect size r_rb_=1). There were 751 (out of 2047) combinations having decoding performance not significantly different from this best performance: 68% of them used both Blocks and Time Between Clicks to predict the local SoA, 15% used Blocks but not Time Between Clicks and 17% used Time Between Clicks but not Blocks.

A Principal Component Analysis (PCA) was applied to reduce data dimensionality and Principal Components (PCs) were ordered in descending order of the variance of the data they explained. We sequentially applied, to an increasing number of PCs, models using Logistic Regression to predict the local SoA. The decoding performance were always significant (p<10^−4^, with an AUC of 0.65 ± 0.10 when including only the first PC, significantly above chance (0.5): Wilcoxon t=132; p<10^−4^; effect size r_rb_=0.96) and reached a ceiling after including the fourth component (AUC = 0.84 ± 0.09, significantly above chance (0.5): Wilcoxon t=0.00; p<10^−4^; effect size r_rb_=1). Together, the first four components explained 66% of the variance. After that, including more components than the first four kept increasing the explained variance but did not significantly increase the decoding performance (p>0.05) and did not differ from the maximum decoding performance obtained when including all components (AUC = 0.85 ± 0.08, p>0.05, **Fig. 4A**). Observing the contributions of the parameters to the components revealed an important weight of the block number for PC4 (**Fig. 4B**), further demonstrating that participants modulate their SoA across the game.

**Figure 4.**
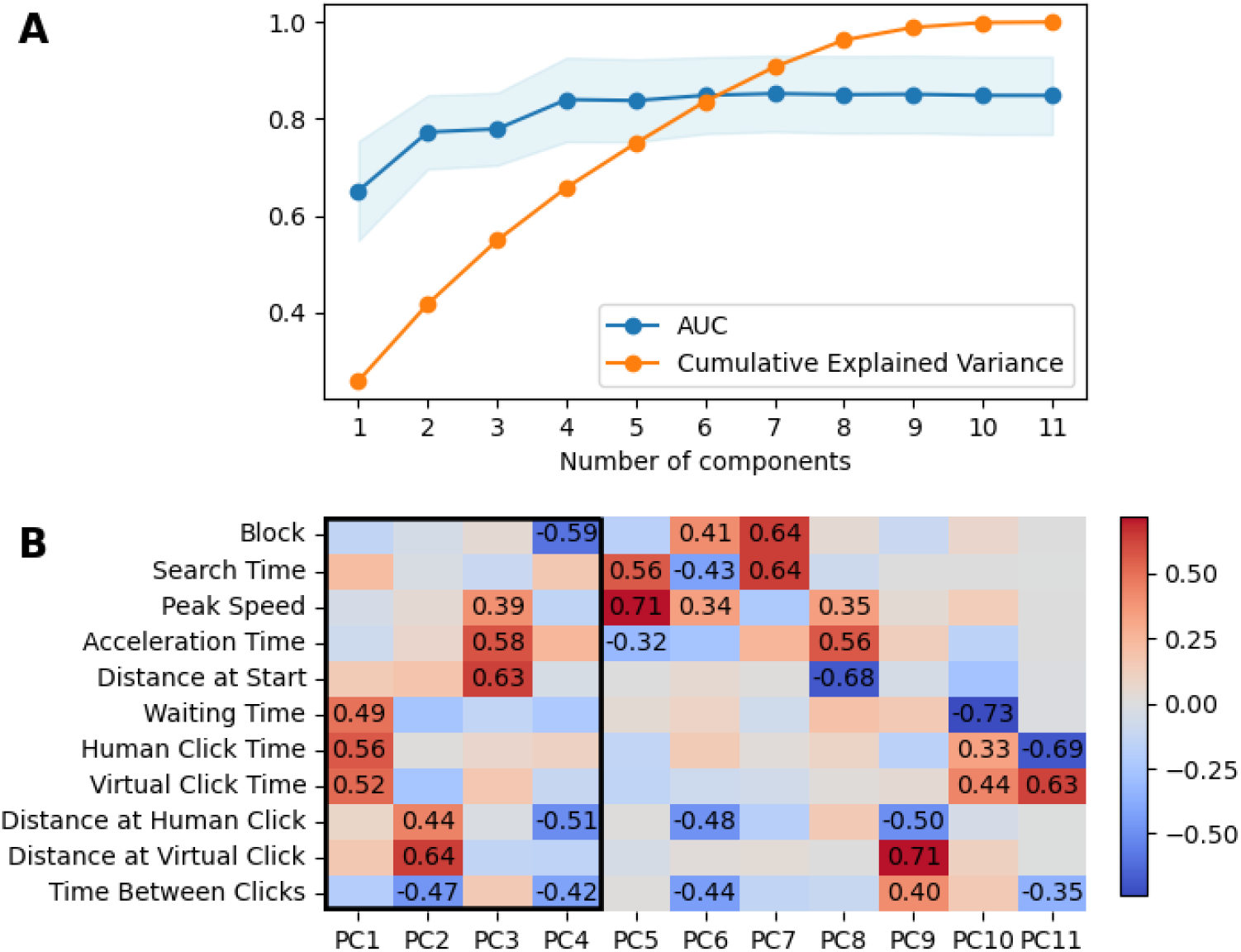
Decoding the local SoA from data features extracted with a Principal Component Analysis (PCA). **A:** Cumulative explained variance calculated by adding the principal components one by one (orange) and decoding performance (Area Under the Curve, AUC) of the logistic regression including an increasing number of components (blue). Four components, explaining 66% of the variance, are sufficient to reach a ceiling decoding performance of 84 ± 9 % (significantly above chance (0.5): Wilcoxon z=0.00; p<10^−4^; effect size r_rb_=1). **B:** Composition of the components. The weights are color-coded; numerical values above 0.3 or below -0.3 are indicated. Note how the block is contributing to the fourth component necessary to reach the ceiling decoding performance.

### Modeling approach: performance and perceived difficulty also influenced the sense of agency

To further test the contribution of these different parameters on the local SoA while minimizing the redundancy in the information they may contain, Ordinary Least Squares (OLS) regression models were fitted to the data to reveal the linear relationship between the dependent variable local SoA and all possible combinations of independent variables measured in each trial. The players were added to the models as random variables, leading to a total of 4094 models (2047 without the players as random variables and 2047 with). The best model was selected as the model with the lowest Akaike Information Criteria (AIC) in order to optimize the goodness of fit of the selected model while privileging its simplicity.

The best model included the players and seven additional explanatory variables of the local SoA (F=195.5; df=115; p<10^−2^; R^2^=0.33, **Fig. 5**, see **Tab 1** for a detail report of coefficients, t-statistics, p-values and confidence intervals for all variables). The intercept of the best model was 0.75, consistent with an average local SoA of 0.74 ± 0.15. Time Between Clicks was the explanatory variable with the highest coefficient (0.20), which confirmed that the local SoA was highly correlated with this variable. We also confirmed the positive relationship between the local SoA and Blocks, independently of the Time Between Clicks (coefficient 0.0086 ± 0.002; p<10-4, Confidence Interval CI=[0.005, 0.012]). The SoA was also positively correlated with the Search Time and the Peak Speed, and negatively correlated with the Distance at Start, at Human Click and at Virtual Click. In other words: the longer they took time to unlock the grid and the faster they were to reach the target, the higher the SoA; the further away they were from the target when the trial started, when they clicked (human click), or when the animation started (virtual click), the lower the SoA. The acceleration time, the waiting time, the human and the virtual click times were not included in the best model, meaning that their contribution, if any, was already included in associated parameters (e.g., the Time Between Clicks is simply a linear combination of human and virtual click times). This result reveals that the participants integrated information about their performance (e.g., how fast they reached the target and how precisely they clicked on it) and about the perceived difficulty (e.g., how far they were from the target when the trial started) to compute their sense of agency.

**Table 1.**
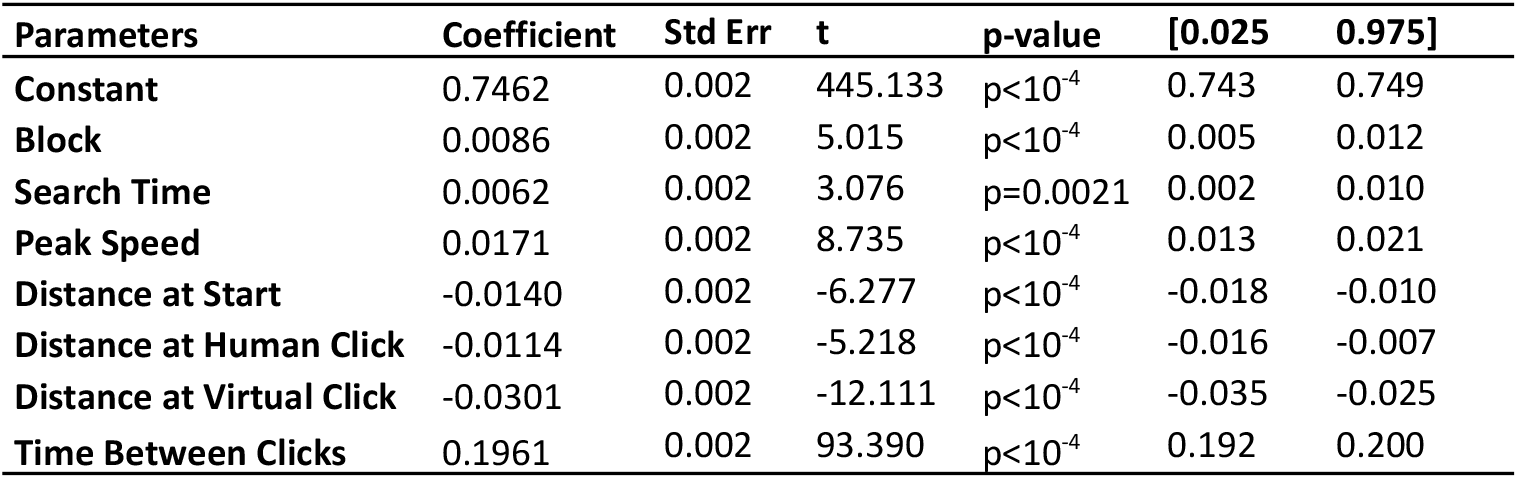
Best model predicting the local SoA’s coefficients and p-value.

**Figure 5.**
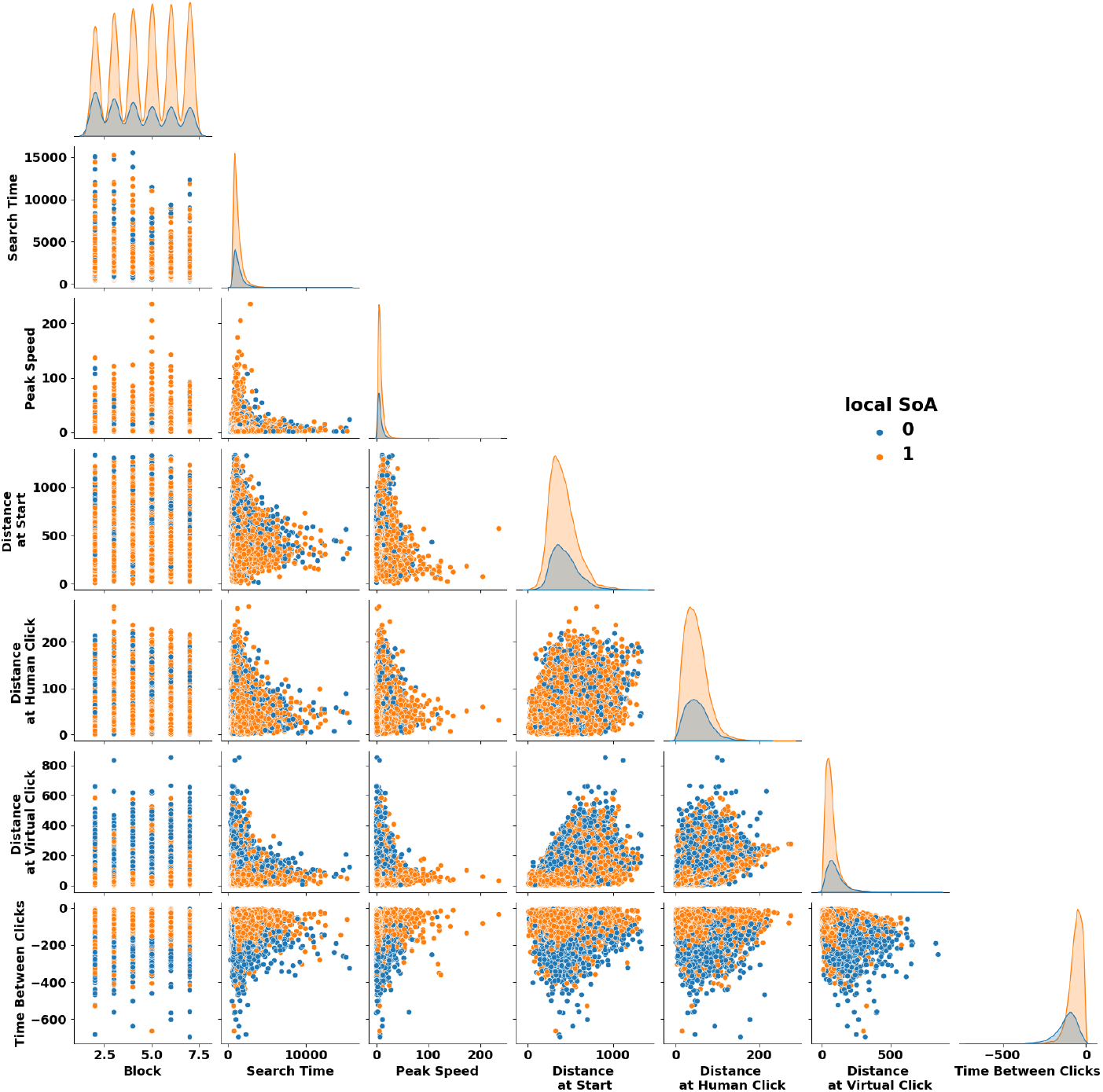
Kernel Density Estimators (KDE) and pairwise relationships between all variables significantly contributing to the local SoA according to the best model (see text for detail). The data is sorted according to the local SoA: blue corresponds to a local SoA of 0 (players believed that the computer won) and orange to a SoA of 1 (players believed that they won). Note how through the course of the game, the number of trials with low local SoA decreased to the benefit of trials with high local SoA (top left) and the distribution of Time Between Clicks for SoA=0 is shifted to the left compared to the distribution for SoA=1 (bottom right). Importantly, all these variables accounted for SoA (independently from each other).

### The sense of agency is computed at multiple timescales

At the end of each block, the players were asked to evaluate their global SoA, i.e., whether they thought they won often against the computer during the last block, using a Likert scale from 1 to 5. Such global SoA was highly correlated to the local SoA (**Supp. Fig. 2B**, R=0.64; p<10^−4^, CI=[0.51, 0.74]) and the shift of corrected global SoA through the game was correlated to the shift of corrected local SoA through the game (i.e., residuals of the correlation between global SoA and Time Between Clicks, and of the correlation between local SoA and Time Between Clicks, were correlated, **Supp. Fig. 2I**, R=0.31; p=0.0009; CI = [0.13, 0.47]). Nevertheless, while the corrected local SoA significantly increased as shown before (**Supp. Fig. 2F**, linear fit coefficient=0.01 ± 0.03; significantly above 0: Wilcoxon t=2240; p=0.0221, effect size r_rb_=0.25), the corrected global SoA did not significantly increase over the course of the game (**Supp. Fig. 2J**, linear fit coefficient=-0.01 +/-0.48; not significantly different from 0: Wilcoxon t=2830; p=0.7302; effect size r_rb_=-0.04). In other words, players increased their SoA on a trial-to-trial basis but did not consciously report a higher SoA at the end of each block. Finally, the corrected local or global SoA shifts were not correlated to the mean local or global SoA, meaning that the ability to shift one’s local or global SoA did not depend on one’s averaged SoA level (all p>0.05; **Supp. Fig. 2D-E, 5G-H**).

At the end of the training and at the end of the game, the players answered 3 open-ended questions to check whether they noticed anything weird, whether they intentionally changed their behavior to trick the game or whether they had any comment. Observation of these comments revealed that a subgroup of players (group 1, N=42) found the game unpleasant: they spontaneously referred to the warnings they received during the game (when they did not correctly play) or to their difficulties, using words such as “penalties” or “frustrating”. Nevertheless, they did not actually receive more warnings than the rest of the players who did not report an unpleasant experience (group 2, N=67, Mann-Whitney: U=1574; p=0.2999). Despite the Time Between Clicks in group 1 being smaller on absolute value than in group 2 (on average, -73 ± 21 ms against -81 ± 20 ms, Mann-Whitney U=1752; p=0.0320; effect size r_rb_=-0.25), the mean global SoA was lower for this group 1 (on average 3.14 ± 0.80 vs. 3.60 ± 0.89, Mann-Whitney U=965; p=0.0058; effect size r_rb_=0.31). Of interest, the shift of the global SoA also significantly differed between groups (Mann-Whitney U=963; p=0.0058; effect size r_rb_=0.32), significantly decreasing in group 1 (significantly below 0: t-test: t(41)=0.02; p=0.0200; Cohen d=0.38) and non-significantly increasing in group 2 (not significantly different from 0: Wilcoxon t=864; p=0.1237; effect size r_rb_=0.22; **Supp. Fig. 3**). On the contrary, the local SoA mean or shift did not differ between the two groups (both p>0.05). Overall, these results suggest that the global SoA estimated at the end of each block is not simply based on the trial-by-trial SoA estimations of this block: it may also integrates other dimensions such as the emotional experience with the game.

### Relationship between low-level (game-related) SoA and high-level (life-related) SoA

At the end of the game, the players answered the Levenson Multidimensional Locus of Control Scales, a questionnaire measuring the belief that what is happening in their own life can be attributed to themselves, to other people having more power, or to chance. Specifically, this scale evaluates the locus of control along these 3 dimensions (I: internal; PO: powerful others; C: chance) in everyday life situations ^26^. Across the group of players, PO and C were correlated (**Supp. Fig. 4**, R=0.61; p<10^−4^; CI=[0.47, 0.71]) but no other correlation was significant, meaning that I and PO/C were, at least partially, capturing independent dimensions. Only the correlation between the mean local SoA and the PO score, and between the mean global SoA and the PO score were significant (R=0.19; uncorrected p=0.0446; CI=[0.00, 0.37] and R=0.21; uncorrected p=0.0299 PO; CI=[0.02, 0.38] respectively) but did not resist correction for multiple comparisons. Thus, no clear link between the SoA mean or shift measured with the game and locus of control in everyday life could be established.

## Discussion

Our online game succeeded in reversing the timeline between action and consequence: it triggered the consequence before the players’ action. This design allowed us to reveal that the sense of agency does not only rely on the objective timeline between actions and consequences, but also on other parameters, such as the performance and perceived difficulty of the game, and, crucially, on the time spent playing the game. We thus confirmed a prediction made by Dennett ^22,23^, that the sense of agency is strongly related to being accustomed to a particular timeline. In line with this prediction, our results demonstrate that just a few tens of minutes are enough to allow adapting the SoA to this unusual situation.

Several important differences with the original “Grey Walter’s precognitive carousel” experiment have to be noted. First, players were not acting spontaneously at their own will time, but they were invited to play fast, i.e., to find the target and, once they found it, to initiate and complete a movement as fast as possible, while trying to maintain accuracy. In this time-constrained situation and contrary to the original experiment, they were not startled to see the animation triggered before they clicked and often failed to notice it. Second, even at the end of the training period, when the animation was triggered obviously early, the players were not excessively surprised to see the animation starting before they had the chance to click on the target, in line with our habits of observing events happening on their own on a computer screen (as compared to, for instance, seeing a light turning on its own, which would be much more surprising). Third, in the experimental blocks, players were told that they played against the computer, so that they expected the animation to sometimes start before their click. Finally, the anticipation was not based on directly reading brain signals related to movement’s preparation, probably attenuating the impression that something read their mind. Nevertheless, the virtual click was not random, but based on the history of the players’ past movements and on the beginning of their current movement. We hypothesize that this association of the virtual click to the players’ own movements is a necessary condition for the fast adaptability of the SoA observed with this game. For the same reasons, we also predict that directly basing the prediction on brain signals will further increase such adaptability, which we will test in a future experiment.

The SoA was also influenced by other factors, including the players’ performance, such as Search Time and Peak Speed (i.e., how fast they were at playing the game) and the Distance at Human Click (i.e., how accurate they were in clicking on the target). This result is consistent with previous studies showing that the SoA is related to performance in various tasks. When action feedback is uncertain, people tend to increase their SoA with increasing performance, even if their performance increase is due to computer assistance, meaning that they actually have less control ^27^. Subjective control ratings have been shown to logarithmically increase as a function of performance ^28^. However, with full automation, the SoA decreases again: people should retain some control (around 10% in some tasks) for optimal SoA and performance ^29^. Some measures have been proposed to capture how judgment of agency is tracking manipulation of controllability beyond judgment of performance ^30,31^. Our results confirm that people integrate their performance to their SoA independently from their true controllability (i.e., how close in time the animation started relative to their button press). Importantly, the fact that players had to make specific movements as fast as possible did not prevent them to increase their SoA, in line with experiments suggesting that action restrictions do not necessarily impact the SoA ^32^.

The SoA was also influenced by how close the cursor was to the target when the animation started (Distance at Virtual Click), which is, with Time Between Clicks, the second factor that fully objective agents would rely on to determine if they won: indeed, if the animation started while they were still far away from the target, then they could not possibly have triggered the animation. Finally, the SoA was negatively correlated with the distance of the cursor to the target at the start of the movement (Distance at Start). We interpreted this factor as belonging to the perceived difficulty of the task: players could feel that if they were far away from the target, it would be difficult to reach it on time, i.e., before the computer. How the computer played was not shared with the players. They were left to imagine an algorithm doing a similar task than the one they were doing (looking for the target and then moving towards it) or doing something entirely different. Somehow, the players thought they were disadvantaged when they had to do a larger effort to reach the target, and this was associated with a decrease of their SoA.

At the end of each Block of trials, players evaluated how often they won, which we called the global SoA, by opposition to the local SoA, which is based on trial-by-trial reports of winning or not. Both local and global SoAs and SoAs’ shifts correlated, however, only the local SoA was sensitive enough to increase with the time spent in the game. Of interest, players seem to integrate other information to derive their global SoA. If they found the game unpleasant, by referring to the warnings they received or their difficulties, their global SoA was lower and decreased through the course of the game. Note that this analysis was not *a priori* planned and derived from examination of players’ free comments. It suggests that multiple scales of SoA exist, relying on different metacognitive processes. A previous study already demonstrated the existence of different neural processing underlying action monitoring and metacognitive evaluation of agency ^33^. Similar investigation should be extended to the hypothetical scales of SoA.

Finally, we found little evidence for a direct relationship between low-level SoA as measured in our game, and a higher-level SoA, a feeling of control over one’s own life and events happening in the world. If any, the SoA, whether local or global, would be higher for players who experience in their life that people other than themselves have more power. This non-intuitive result will need to be investigated with experiences specifically designed for this purpose.

In this article, we introduced a novel game, which is able to trigger a consequence of an action shortly before players perform the action. We demonstrated that players learnt, despite the unusual timings, that they were at the origin of the animation and adapted their SoA accordingly in only a few tens of minutes. We showed robust evidence that humans can adapt their sense of agency to such a very unusual situation, that will nevertheless become more common as AI is becoming more pervasive.

## Acknowledgment

This work was supported by the Impulsion Grant from IDEXLYON (Initiative d’Excellence de l’Université de Lyon) and ANR MyAct to Marine Vernet.

The authors would like to thank Sonia Alouche, Florence Tamissa-Leger, Célia Farge, Lydie Merle-Brigaud and Lesly Fornoni for administrative support.

This article is dedicated to the memory of Daniel Dennett (1942-2024).

## Material and methods

### Participants (inclusion and ethics)

One hundred and twelve naive healthy adult participants were included in this study (55 females, 14 left-handed and 2 ambidextrous by self-reports, 27 ± 7 years old). The participants reported no history of neurological or psychiatric conditions, except one participant who reported an ADHD diagnosis in the free comments and was excluded from further analyses. Two other participants were excluded: one always reported being the agent and the second one only reported twice that she was not the agent (over 480 trials). All remaining 109 participants (53 females, 13 left-handed, 2 ambidextrous, 27 ± 7 years old) reported that they were the agent in 371 ± 7 trials (range [146, 473]) and were included in all analyses. The participants were recruited via mailing lists or a recruitment platform (Prolific) and received monetary compensation for their participation. The study was approved by an ethical committee (Inserm CEEI, IRB00003888, IORG0003254, FWA00005831, Opinion # 22-928, ADAPT-AGENT). The experiment has been designed in 2021 and the data collected in 2022.

### Tasks

Participants performed an online game, written in JavaScript and available through Pavlovia, which was based on a visual search task. In each trial, players had to find a visual target in a 7-by-4 grid of distractors with abstract shapes **(Supp. Fig. 5A)**. The target had a different shape and color than all the distractors and was situated between two and five times its size from the cursor at the start of the trial. Once the target was found, players unlocked the grid by pressing the spacebar and moved the cursor with a mouse to click on the target, which triggered a short animation. The animation consisted in a series of effects: the target flashing and pulsating while the entire grid (except the target) shook and progressively vanished (all these effects lasting up to 160 ms) and a circle appearing around the target with black particles coming out of it (as if the target was exploding) for 567 ms. Players were not allowed to move the mouse during the research time before pressing the spacebar and once the spacebar was pressed, they had to immediately perform a fluid and direct movement towards the target, otherwise, the participant would get a warning message and the trial would start over (for a list of potential warnings, see **Supp. Fig. 6**). The experiment started with a short habitation block, which ended once the participant performed 5 trials in a row without any warning. The experiment then continued with one training block and 6 experimental blocks, each made of 80 correct trials (without warning). The training block was used to further train the participant to make consistent hence predictable movements and to train an algorithm to predict the participant’s click time.

The prediction, which was updated on each trial through the course of the experiment, was based on minimizing the following equation:

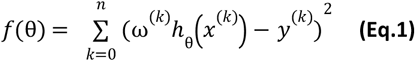

where *k* = 0 is the first trial and *k* = *n* is the last trial; *y*^(*k*)^ is the time between the spacebar press and the players’ click on the target on trial *k*; *x*^(*k*)^ is the input vector on trial *k*, with four parameters: Waiting Time (i.e., time between the spacebar press and the beginning of the cursor movement, when its speed exceeded 0.2 pixels/ms), Distance at Start (i.e., distance between the cursor and the target at the beginning of the cursor movement), Peak Speed (i.e., last speed recorded before not registering any speed increase for 2 samples in a row i.e. for 32 ms) and Acceleration Time (i.e., time between the beginning of the cursor movement and the time it reached its peak speed); 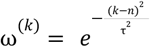 is the weight attributed to the trial *k* (recent trials receiving more weights than earlier ones, with τ = 50); and *h* (*x*) = 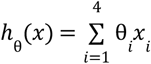 is the dot product between the input vector *x* and a vector θ, whose components are determined by minimizing the function in Eq.1, which was done by solving matrix operations. Once θ was updated for each trial, predicting the click time was then made by estimating *y* = *h* _θ_ (*x*) for the current trial. The actual “virtual click” time, corresponding to the triggering of the animation by the algorithm during the main experiment, was set to occur before the predicted “human click” by subtracting 2 ^*^ *mean*(*weighted prediction error*) + *std*(*weighted prediction error*) from the predicted click time (the weights being described by ω^(*k*)^ used in Eq. 1). In addition, if the value of *A* = *mean*(*weighted prediction error*) + *std*(*weighted prediction error*) was below a threshold of 100, a number randomly chosen from a uniform distribution between 0 and 100 *− A* was also subtracted, to ensure that the click occurred sufficiently early relative to the human click **(Supp. Fig. 7)**.

In the last 10 trials of the training block, the animation was programmed to start even earlier (in the first 5 trials, it was triggered at the predicted time of the click minus an increasing multiple of 40 ms; and for the next 5 trials it was triggered as soon as the cursor’s movement start was detected). The goal was to make the start of the animation obviously before the player’s click. It was indeed triggered rapidly after the spacebar (on average 337 ± 192 ms) so that the players either clicked after the animation started (on average 192 ± 116 ms after, 49% of the trials) or did not even have the time to click (51% of the trials). At the end of this block, the players were asked to comment freely on whether they noticed anything in the last few trials. Most players (90; 83%) noticed that the animation started before they clicked; 8 other players reported that something changed, such as the game or their movements becoming faster, or the target being highlighted when they pressed the spacebar; 11 players did not report anything strange. Then, all the players were informed that the computer “clicked” on the target before they could do it and were instructed new rules for the following blocks. They should now answer after each trial if they thought they clicked on the target first (i.e., they felt the agent, SoA = 1) or if it was the computer (i.e., they didn’t feel the agent, SoA = 0, **Supp. Fig. 5B**). A new warning message was also introduced to avoid a situation in which participants would intentionally let the computer win by purposely delaying their movement (see **Supp. Fig. 6)**. During the experimental blocks, the algorithm was successful at performing a “virtual click” shortly before the participant‘s click in 86 ± 11 % of the trials without warnings, with an average anticipation of 78 ± 21 ms

(corresponding to 406 ± 238 ms after the players pressed the spacebar). The duration of the 6 experimental blocks was 41 ± 12 min and the total duration of the game was 54 ± 16 min).

To keep the players engaged with the game, blocks were called “levels” **(Supp. Fig. 5A&E)**. The shapes became more complex in higher levels, but the target was always identifiable based on its unique color in the grid (note that the color randomly changed every trial). Every 20 trials, a performance review was delivered, including the Search Time, the number of warnings, and how these metrics compared to preceding performance reviews (**Supp. Fig. 5D**, see also **Supp. Fig. 1** for an evolution of the parameters through the course of the game). At the end of each experimental block, the players had to estimate with a Likert scale from 1 to 5, whether 1) they thought they won more often than the computer and 2) they thought they were accurate in their assessment of the first question **(Supp. Fig. 5E)**.

At the end of the experiment, the players were asked whether they tried to “trick” the computer or intentionally change their behavior at any point during the experiment, as well as if they had comments about the task. Then, they answered the Levenson Multidimensional Locus of Control Scales ^26^, a questionnaire evaluating the locus of control along 3 dimensions (internal, powerful others and chance), in which the players rated for each of 24 items how much they agree with the statement on a scale from -3 (strongly disagree) to +3 (strongly agree) without a 0 option. The entire experiment timeline can be seen in **Supp. Fig. 8**.

### Statistical analyses

Shapiro tests were used to check the normality of the data distributions. When data was normal, testing against zero was performed with t-tests (effect size was calculated by Cohen’s d). When data was not normal,, Wilcoxon tests were used for testing against zero (or against chance at 0.5 for the decoding analysis) and Mann-Whitney U tests were used for comparing two distributions (for both, effect size was calculated by rank biserial correlation r_rb_). For correlations, Pearson coefficients were used (95% confidence intervals were computed when the coefficients were computed at the group level; computations at the participant level were followed by the statistics described above). All tests were two-sided and a cut-off point of 0.05 for the p-values was used.

### Data and Code availability

Data supporting the findings and code used in this study is publicly available at https://github.com/marinevernet/adapt-agent.git

---

**Supplementary Figure 1.**
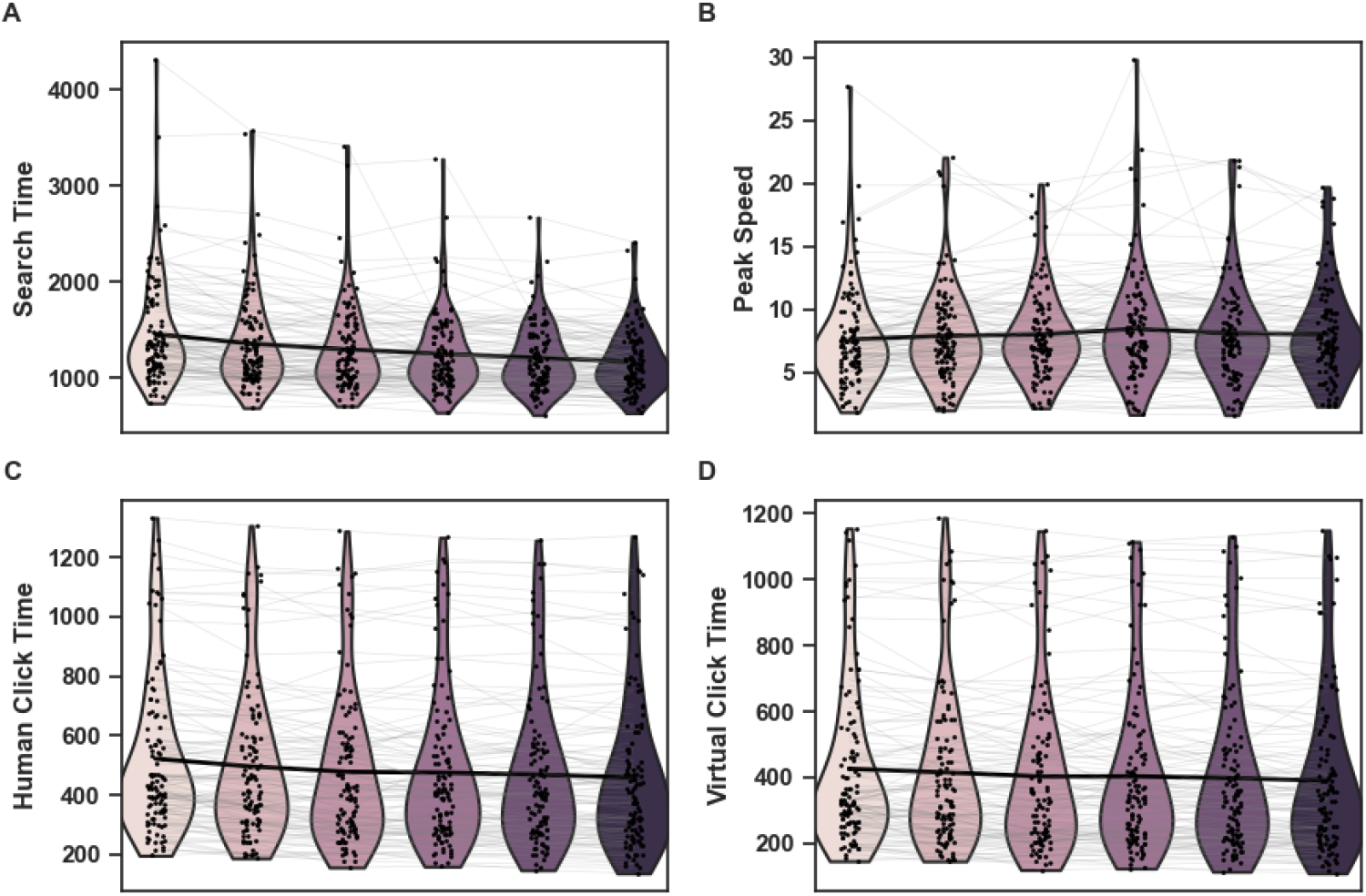
Significant evolution of players’ performance and of game’s click time. **A. Search time decreased** (linear fit coefficient=-54.69 ± 82.27; significantly below 0: Wilcoxon t = 193; p<10^−4^; effect size r_rb_=-0.94). **B. Peak Speed increased** (linear fit coefficient=0.09 ± 0.38; significantly above 0: Wilcoxon t = 1848; p = 0.0005; effect size r_rb_=0.38). **C. Human Click Time decreased** (linear fit coefficient=-11.43 ± 16.14; significantly below 0: t-test t(108)=-7.36; p<10^−4^; effect size d=0.71). **D. Virtual Click Time decreased** (linear fit coefficient=-6.58 ± 15.43; significantly below 0: Wilcoxon t = 1442; p<10^−4^; effect size r_rb_=-0.52). Note that Waiting Time, Acceleration time, Distance at Start and Distance at Virtual Click did not evolve through the course of the game (all p>0.05, not shown). Distance at Human Click increased (linear fit coefficient=0.33 ± 1.46; significantly above 0: Wilcoxon t = 2226; p = 0.0198; effect size r_rb_=0.26) but this increase did not resist correction for multiple comparisons.

**Supplementary Figure 2.**
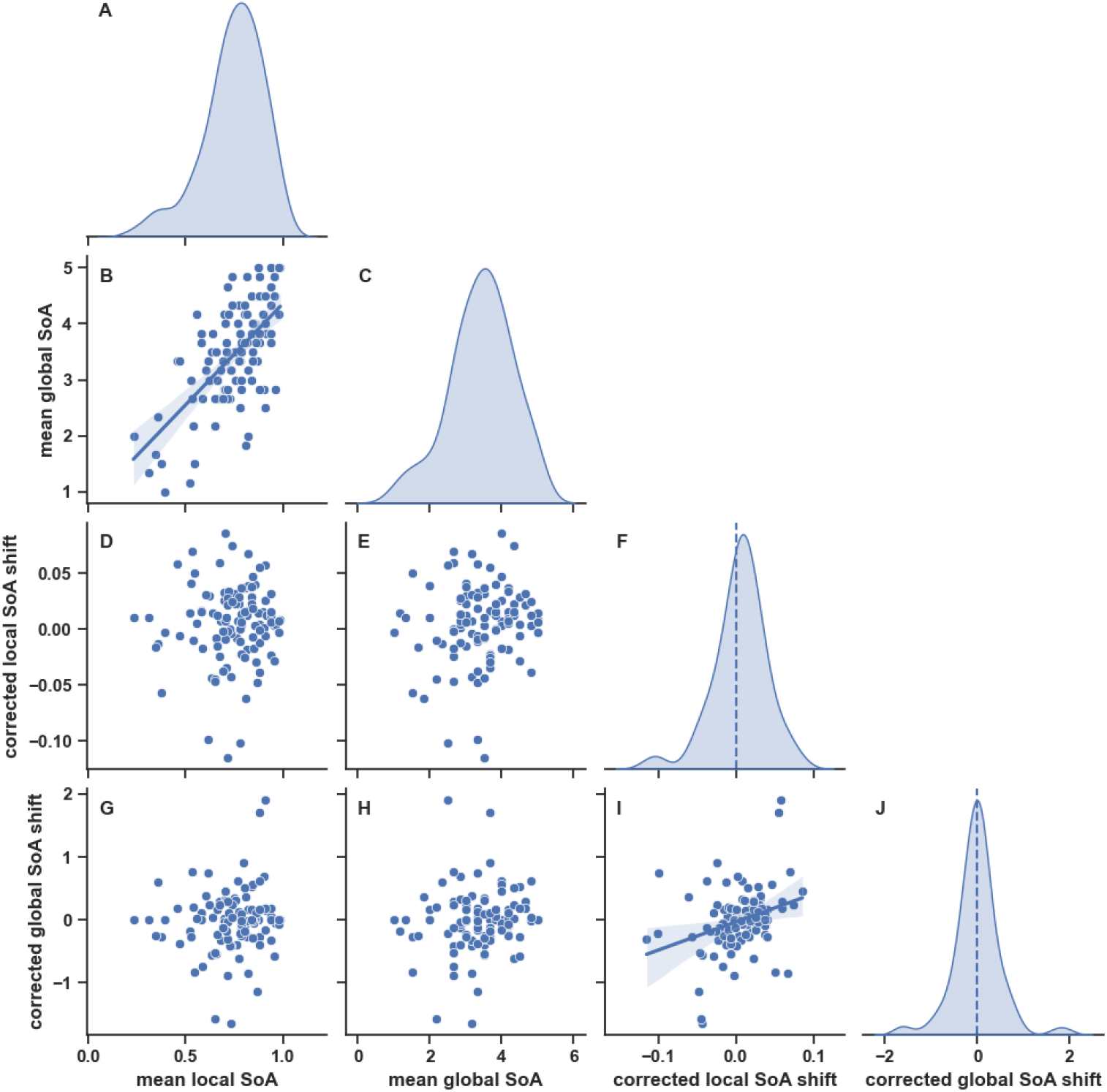
Kernel Density Estimators (KDE) and pairwise relationships between mean global and local SoA and corrected local and global SoA shifts. **A, C, F & J:** Distribution of the values (KDE) for the four variables of interest. Note the distribution of the corrected local SoA shift significantly above 0 (0.01 ± 0.03; Wilcoxon t=2240; p=0.0221, effect size r_rb_=0.25, **F**), meaning that the local SoA increased through the course of the game (independently from the Time Between Clicks tested). **B, D, E, G, H:** Pairwise relationship between the four variables of interest. Note the significant correlation between mean local and global SoA (R=0.64; p<10^−4^; CI = [0.51, 0.74], **B**) and between the corrected local and global SoA shifts (R=0.31; p=0.0009; CI = [0.13, 0.47], **I**).

**Supplementary Figure 3.**
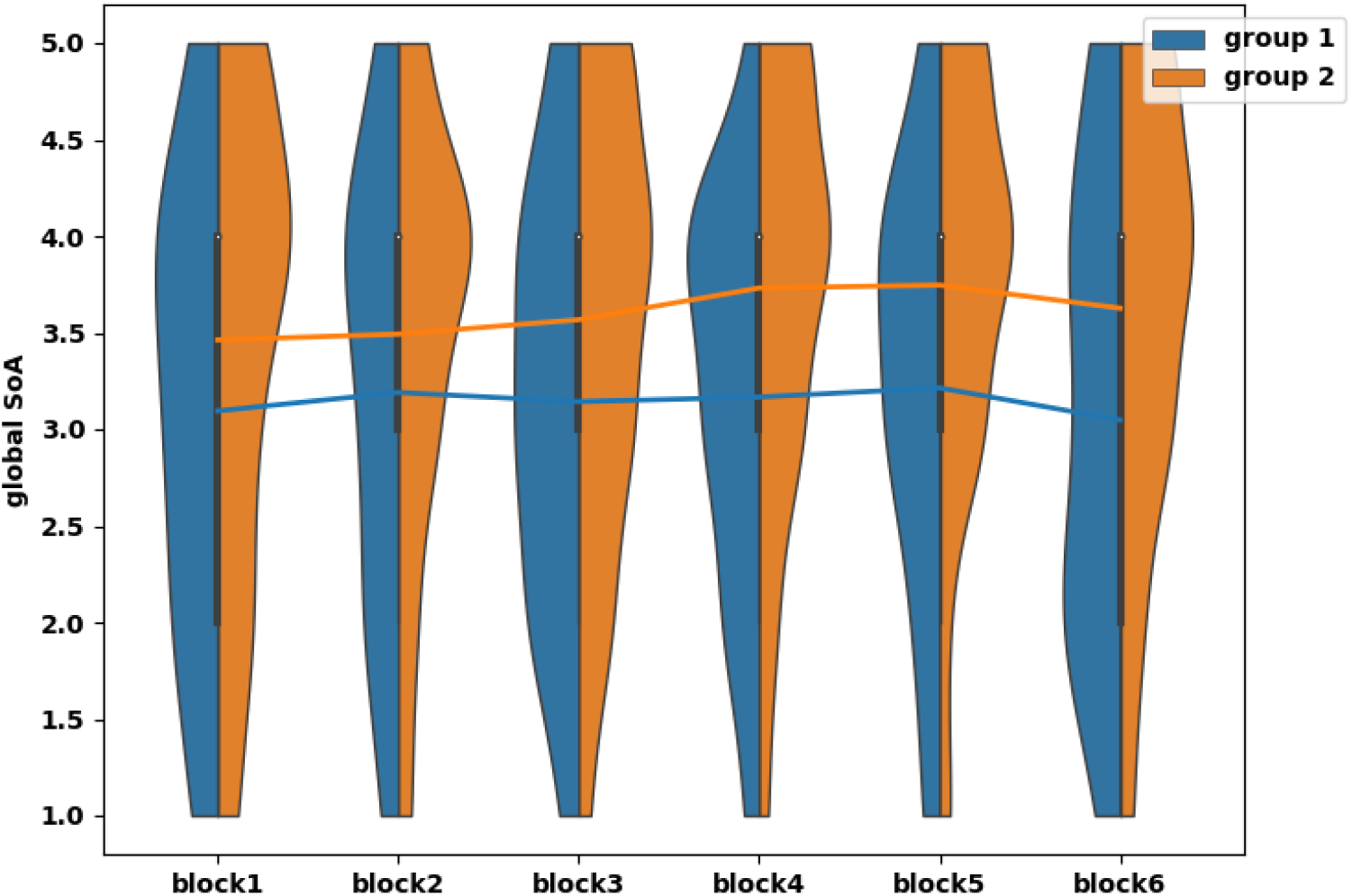
Evolution of the global SoA through the course of the game separately for players who mentioned an unpleasant experience with the game (group 1, N=42) and the rest of the players who did not (group2, N=67). Note the lower mean global SoA for group 1 than group 2 (Mann-Whitney U=965; p=0.0058; effect size r_rb_=0.31) and the global SoA significantly decreasing only in group 1 (t-test: t(41)=0.02; p=0.0200; Cohen d=0.38) while non-significantly increasing in group 2 (Wilcoxon t=864; p=0.1237; effect size r_rb_=0.22).

**Supplementary Figure 4.**
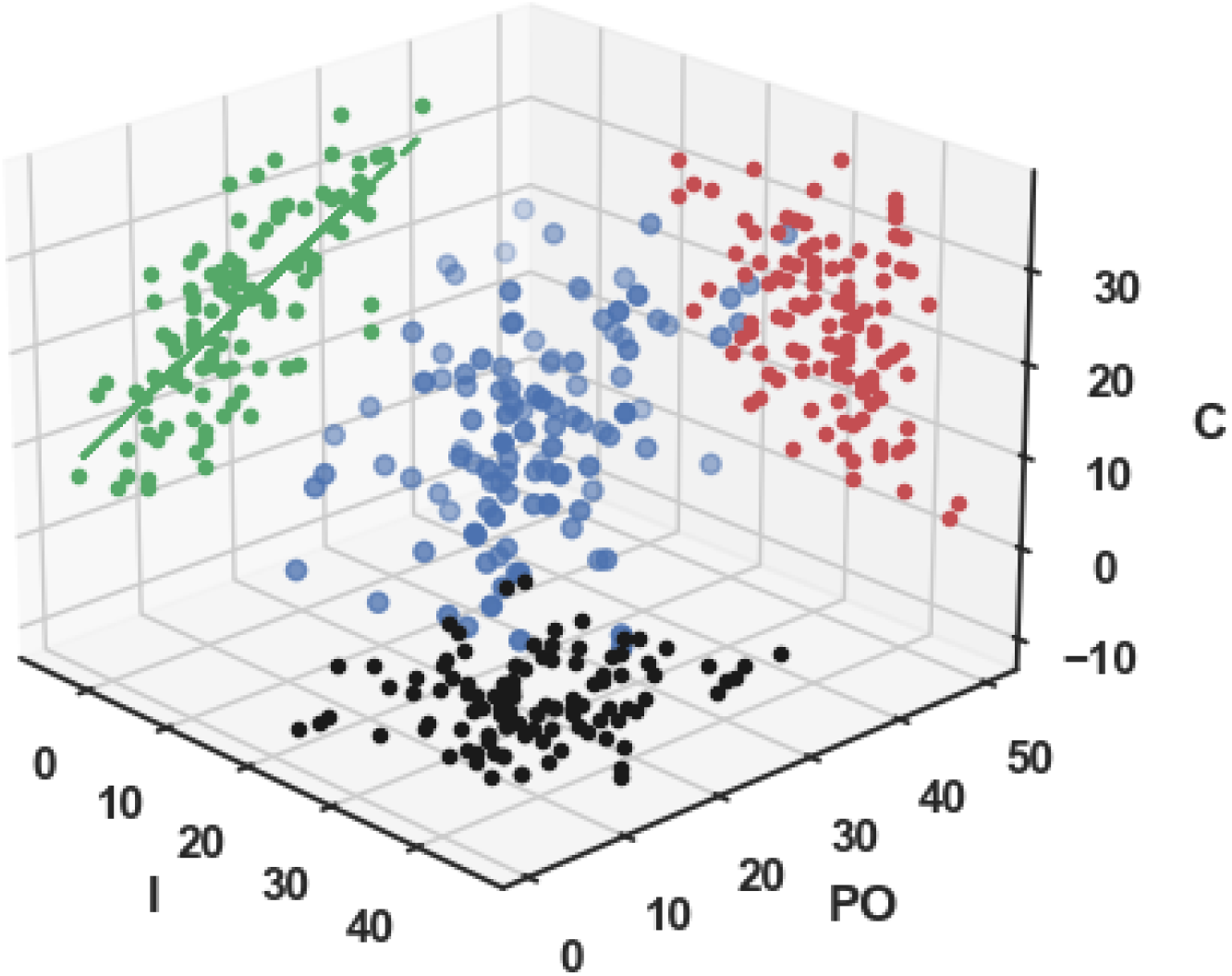
Ratings of the Locus of Control along 3 dimensions: Internal (I), Powerful Others (PO) and Chance (C). Note the significant correlation between PO and C, R=0.61; p<10^−4^; CI=[0.47, 0.71])

**Supplementary Figure 5.**
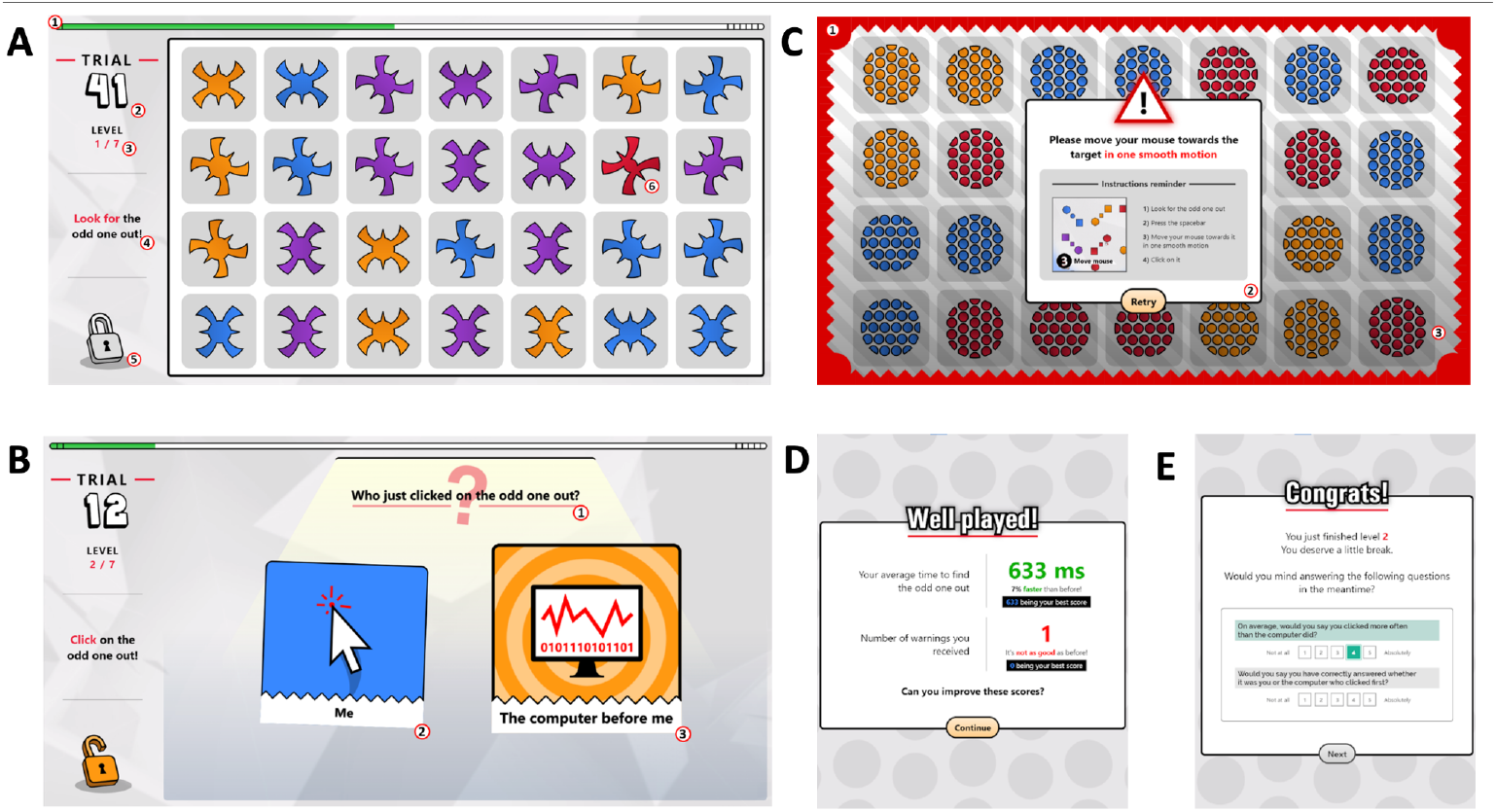
Main Screenshots from the game. **A. Visual search grid.** 1. Progress bar. 2. Trial indicator. 3. Block (=level) indicator. 4. Instruction. 5. Locked/Unlocked grid Indicator 6. Target. **B. Response screen**. 1. Question. 2. Answer “me” (local SoA=1). 3. Answer “computer” (SoA=0). Note that the cartoon for the answer “computer” is currently bigger because it is selected by a player for this particular trial. **C. Warning**. 1. Red border. 2. Pop-up box. 3. Animation corresponding to a correct sequence of events. **D. Feedback on Performance**. Delivered every 20 trials. **E. Subjective ratings of global SoA and confidence**. At the end of each block.

**Supplementary Figure 6.**
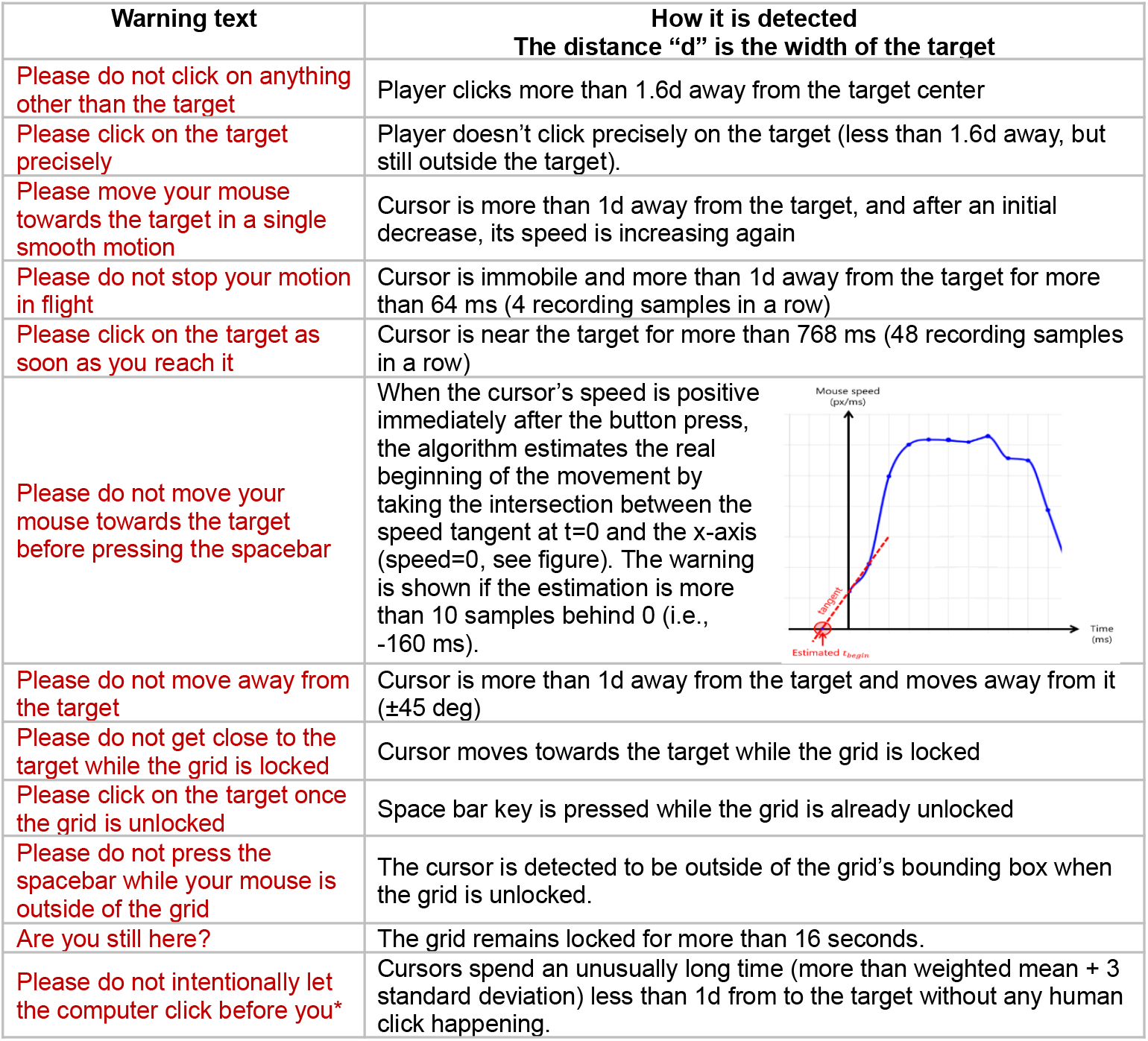
Possible warnings and their causes. *Note that the last warning was only introduced for the experimental blocks, after the training.

**Supplementary Figure 7.**
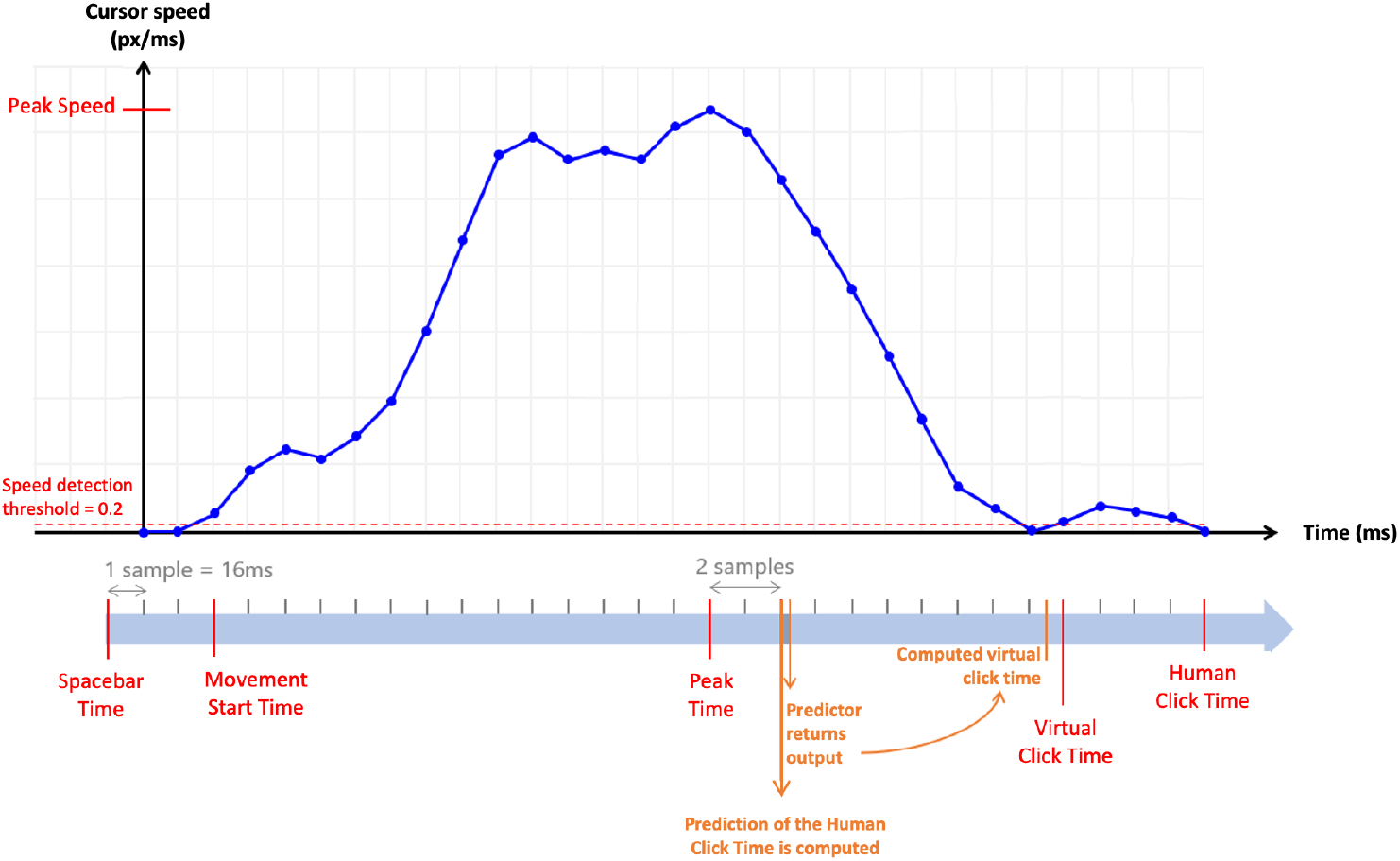
Timeline of a typical trial. As soon as the peak speed is detected, the following 4 parameters are fed to the the algorithm computing the virtual click time: Search Time (Spacebar Time - Trial Start Time), Distance at Start (i.e., distance between the cursor and the target at the Movement Start Time), Peak Speed (i.e., last speed recorded before not registering any speed increase for 2 samples in a row, i.e. for 32 ms) and Acceleration Time (Peak Time - Movement Start Time). Once computed, the actual Virtual Click will occur at the next sample. Note that if the computed virtual click time is before the current time, it is triggered right away. In addition, if the cursor movement is too erratic around the target, i.e., if there are more acceleration and deceleration phases in the current trial than in any of the preceding trials, the virtual click is canceled.

**Supplementary Figure 8.**
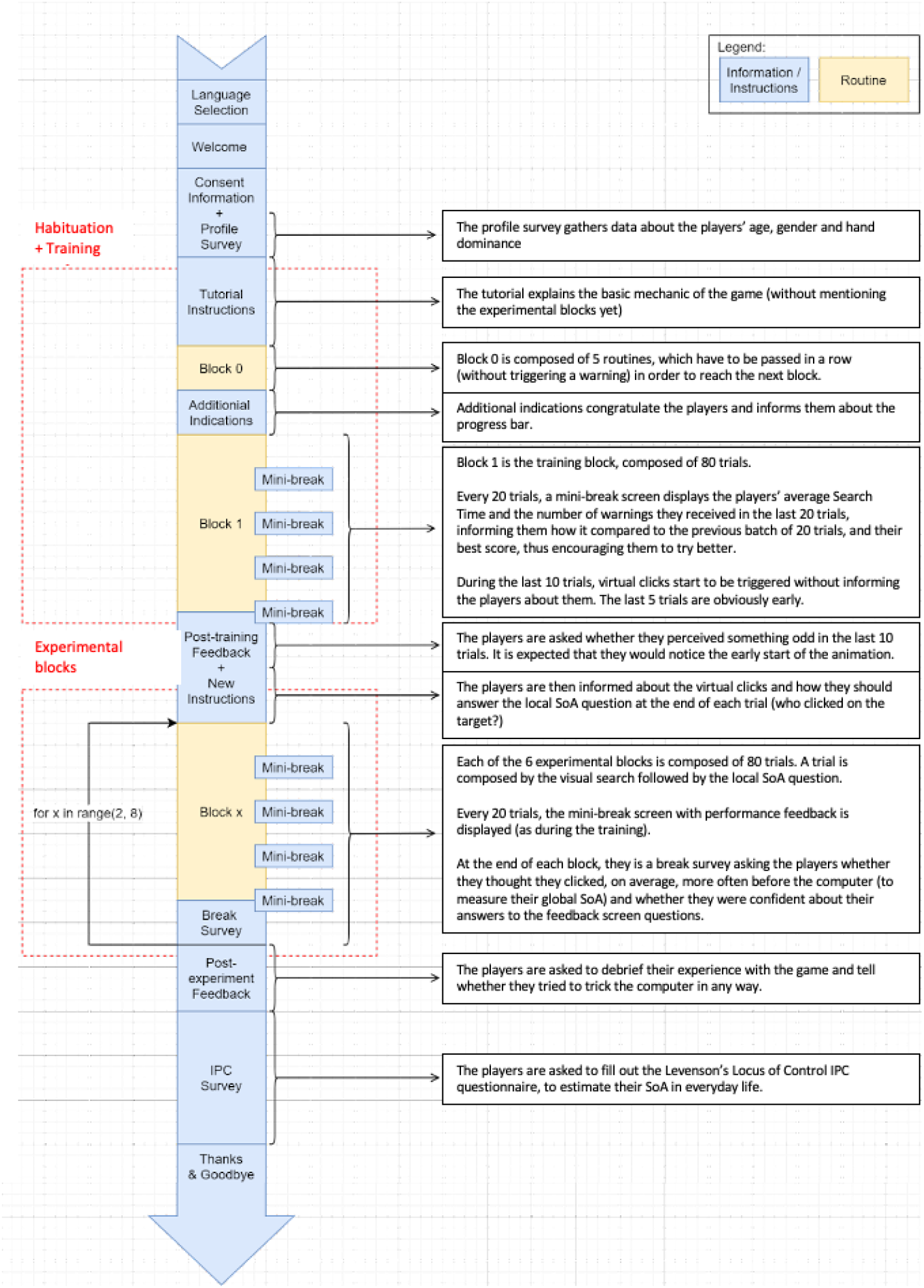
Timeline of the entire experiment.

